# Increasing proteome depth while maintaining quantitative precision in short gradient data-independent acquisition proteomics

**DOI:** 10.1101/2022.09.12.507556

**Authors:** Joerg Doellinger, Christian Blumenscheit, Andy Schneider, Peter Lasch

## Abstract

The combination of short liquid chromatography (LC) gradients and data independent acquisition (DIA) by mass spectrometry (MS) has proven its huge potential for high-throughput proteomics. This methodology benefits from the speed of the latest generation of mass spectrometers, which enable short MS cycle times needed to provide sufficient sampling of sharp LC peaks. However, the optimization of isolation window schemes resulting in a certain number of data points per peak (DPPP) is understudied, although it is one of the most important parameters for the outcome of this methodology. In this study, we show that substantially reducing the number of DPPP for short gradient DIA massively increases protein identifications while maintaining quantitative precision. A deeper analysis of the underlying effects shows that a reduction of DPPP increases the selectivity of the data analysis, allowing fragment ions to be tracked over a longer period of the LC gradient. This in turn increases the number of precursors identified per protein, keeping the number of data points per protein nearly constant even at long cycle times. When proteins are inferred from its precursors, quantitative precision is maintained at low DPPP while greatly increasing proteomic depth. This strategy enabled us quantifying 6018 HeLa proteins (> 80,000 precursor identifications) with coefficients of variation below 20% in 30 min using a Q Exactive Orbitrap mass spectrometer, which corresponds to a throughput of 29 samples per day. This indicates that the potential of high-throughput DIA-MS has not been fully exploited yet.

## INTRODUCTION

Technical progress has transformed mass spectrometry (MS)-based proteomics into a high-throughput technology for analysis of large sample cohorts. This contributes to gain new insights in many different areas, including precision medicine, cell biology, biomarker research and single cell analysis. Many impactful approaches rely on the use of short gradients combined with data-independent acquisition (DIA) and AI-supported data analysis ^1-3^. This strategy benefits from the increased speed and sensitivity of the latest generation mass spectrometers. However, due to the high costs of these instruments and fast development of new hardware, which requires large investments every few years to keep the equipment of proteomic labs state-of-the-art, the spread of this technology often falls behind topical demands for high-throughput proteomics.

Research in improving DIA-MS has focused on the implementation of the DIA acquisition strategy on new MS hardware ^2, 4^, the development of novel acquisition schemes ^3, 5, 6, 7, 8^ and the improvement of data analysis ^9, 10^ as well as library generation ^11, 10, 12, 13^. However, one of the key aspects of setting up of DIA acquisition methods is understudied, namely determining the optimal number of data points per peak (DPPP) ^14^. Usually, a number of DPPP is chosen based on rule of thumbs or personal experience rather than experimental data, which in turn specifies a certain cycle time from which a corresponding number of isolation windows follows. In general, this strategy favors the use of fast scanning mass spectrometers enabling the use of more windows at a given cycle time and so the identification of more proteins. It is general knowledge, that lowering the number of DPPP results in decreased quantitative accuracy and precision. In this study we show, that the concept of designing DIA acquisition schemes based on a certain number of data points per peak has limitations when considering the fact, that the majority of proteomic studies report protein level data based on precursor measurements. Our data prove, that isolation window optimizationcan increase the number of identified precursors so massively, that quantitative precision of proteins is maintained at a largely improved proteome depth even for low numbers of DPPP. This strategy enabled us to quantify 6018 HeLa proteins with coefficients of variations below 20% using a Q Exactive HF orbitrap mass spectrometer at rather moderate scanning speed (12 Hz) in 30 min LC gradients. In total, 7318 proteins were identified in this triplicate analysis from an *in silico* predicted human library. These results correspond to a throughput of 29 samples per day and were achieved using a regular nanoLC and a mass spectrometer that came on the market more than 7 years ago. Noteworthy, our results are quite on pair with published data using the latest generations of MS instruments ^1, 2^, which shows, that the potential of high-throughput DIA-MS has not been fully exploited yet and that older MS instruments are also well suited for proteomic studies using short gradients.

## EXPERIMENTAL PROCEDURES

Cultivation. *Escherichia coli* K-12 (DSM 3871) was cultivated on Tryptic Soy Agar (TSA) ReadyPlates™ (Merck, Darmstadt, Germany) at 37°C overnight. *Saccharomyces cerevisiae* strain S288C (ATCC 204508) was cultivated on MT agar plates supplemented with hemoglobin and charcoal (MTKH) for 48 h at 37°C. Cells were harvested using an inoculating loop and washed in 2 × 1 mL phosphate-buffered saline (PBS) for 5 min at 4,000 × g and 4°C. HeLa cells (ATCC® CCL-2™) were cultivated in Dulbecco’s Modified Eagle’s Medium (DMEM) supplemented with 10% fetal calf serum (FCS) and 2 mM L–glutamine at 37°C and harvested at 90% confluency by scraping. Cells were washed in 2 × 2 mL PBS for 5 min at 400 × g and 4°C. Cell numbers in aliquots of the samples were determined using Neubauer counting chambers. For each sample cell numbers were determined in triplicates.

Sample preparation. *E. coli, S. cerevisiae* and HeLa cells were prepared for proteomics using *Sample Preparation by Easy Extraction and Digestion (SPEED)* ^15^. At first cells were resuspended in trifluoroacetic acid (TFA) (Optima™ LC-MS grade, Thermo Fisher Scientific, Waltham, MA, USA) (sample/TFA 1:10 (v/v)) and incubated at room temperature for 3 min. Samples were neutralized with 2M TrisBase using 10 × volume of TFA and further incubated at 95°C for 5 min after adding tris(2-carboxyethyl)phosphine (TCEP) to a final concentration of 10 mM and 2-chloroacetamide (CAA) to a final concentration of 40 mM. Protein concentrations were determined by turbidity measurements at 360 nm using a NanoPhotometer® NP80 (Implen, Munich, Germany), adjusted to 1 µg/µL using a 10:1 (v/v) mixture of 2M TrisBase and TFA and then diluted 1:5 with water. Digestion was carried out for 20 h at 37°C using Trypsin Gold, mass spectrometry grade (Promega, Fitchburg, WI, USA) at a protein/enzyme ratio of 100:1. Resulting peptides were desalted using Pierce™ peptide desalting spin columns (Thermo Fisher Scientific) according to manufacturer’s instructions and concentrated using a vacuum concentrator. Dried peptides were suspended in 20 µL 0.1% TFA and quantified by measuring the absorbance at 280 nm using the NanoPhotometer® NP80 (Implen). Peptide mixtures of different species were prepared from SPEED preparations of *E. coli* and HeLa cells as well as of a commercially available yeast protein digest (Promega), The ratios of the peptide amount of each species within the three mixtures are given in Table S1.

Liquid chromatography and mass spectrometry. Peptides were analyzed on an EASY-nanoLC 1200 (Thermo Fisher Scientific) coupled online to a Q Exactive™ HF mass spectrometer (Thermo Fisher Scientific). 1 µg of peptides were separated on a PepSep column (15 cm length, 75 µm i.d., 1.5 µm C18 beads, PepSep, Marslev, Denmark) using a stepped 30 min gradient of 80% acetonitrile (solvent B) in 0.1% formic acid (solvent A) at 300 nL/min flow rate: 4–9% B in 2:17 min, 9-26% B in 18:28 min, 26–31% B in 3:04 min, 31–38% B in 2:41 min, 39–95% B in 0:10 min, 95% B for 2:20 min, 95–0% B in 0:10 min and 0% B for 0:50 min. Column temperature was kept at 50°C using a butterfly heater (Phoenix S&T, Chester, PA, USA). The Q Exactive™ HF was operated in a data-independent (DIA) manner in the m/z range of 345–1,650. Full scan spectra were recorded with a resolution of 120,000 using an automatic gain control (AGC) target value of 3 × 10^6^ with a maximum injection time of 100 ms. The full scans were followed by various numbers of DIA scans (Tables S2-8). In order to introduce retention time dependent segments of DIA cycles with fixed cycle times but varying window widths, the window centers are deposited in the inclusion list of the QE method editor along with start and end times. Furthermore, the runtime and the window width of each single DIA scan event needs to be harmonized in accordance to the inclusion list (FIGURE S1). DIA spectra were recorded at a resolution of 30,000 using an AGC target value of 3 × 10^6^ with the maximum injection time set to auto and a first fixed mass of 200 Th. Normalized collision energy (NCE) was set to 27% and default charge state was set to 3. Peptides were ionized using electrospray with a stainless-steel emitter, I.D. 30 µm (PepSep) at a spray voltage of 2.1 kV and a heated capillary temperature of 275°C.

Data analysis. Protein sequences of *Homo sapiens* (UP000005640, downloaded 24/11/21), *E. coli* K-12 (UP000000625, downloaded 26/11/21), and *S. cerevisiae* strain ATCC 204508 (UP000002311, downloaded 29/11/21) were obtained from UniProt. Spectral libraries were predicted using the deep-learning algorithm implemented in DIA-NN (version 1.8) ^9^ with strict trypsin specificity (KR not P) allowing up to one missed cleavage site in the m/z range of 300 – 1,800 with charges states of 1 – 4 for all peptides consisting of 7-30 amino acids with enabled N-terminal methionine excision and cysteine carbamidomethylation. The mass spectra were analyzed in DIA-NN (version 1.8) using default settings including a false discovery rate (FDR) of 1% for precursor identifications with enabled “match between run” (MBR) option for technical triplicates. The resulting precursor.tsv and pg_matrix.tsv (Lib.Q.Value = 1%) files were used for further analysis in R (version 4.1.3) Perseus (version 1.6.5.) ^16^. Please note, that the numbers of identified protein groups, which were extracted from the pg_matrix.tsv, differs from the numbers reported in the stats.tsv file as DIA-NN uses different q-values for filtering in these two files. Differentially abundant proteins of the species mixture samples were identified using FDR-adjusted p-values from a t-test with a permutation-based FDR of 0.05 and s0 = 0.1 after normalization of the log-2 transformed MaxLFQ intensities using row cluster subtraction of the human proteins.

## RESULTS AND DISCUSSION

### DIA window optimization

To achieve a balance between sample throughput and effective use of MS scan time, a 30 min LC gradient was used in this study. This corresponds to a throughput of ∼ 29 samples per day as the actual duration of one run on this nanoLC setup is ∼ 50 min. The peptide elution window is ∼ 26 min and so the mass spectrometer is measuring peptides for ∼ 50% of the actual run time. A segmented gradient was used to achieve a uniform elution of peptides within the elution window. At first, we aimed to determine the optimal number of data points per peak (DPPP) with respect to protein identification and quantification. HeLa samples were used to determine the retention time (rt) and mass to charge (m/z) distribution of all detectable precursors from measurements using narrow isolation windows (4 m/z widths with 2 m/z overlap) and gas-phase fractionation (GPF) (8 × 100 m/z, 350 – 1,150 m/z). For this purpose, a library of all possible human precursors was predicted using Prosit ^11^. The number of isolation windows for selected numbers of DPPP (1, 1.25 and 1.5) were calculated from the average chromatographic peak width (full width at half maximum, FWHM) of the precursors identified in the GPF data using DIA-NN and the durations of the MS^1^ and DIA scans (TABLE S3). The dynamic widths of the isolation windows were then selected such as the number of precursors identified in the GPF data was kept constant for each window ^14^. It should the noted, that the effective numbers of DPPP in the single-run experiments reported by DIA-NN were slightly above the calculated values, because the sensitivity in the GPF experiment exceeds the single-shot experiments (TABLE S3). The DIA acquisition strategy with staggered windows and forbidden zones described by Pino et al., was selected as a reference method ^17^. Therefore, the number of isolation windows was calculated from the chromatographic peak widths (6σ) of the precursors identified in the GPF data determined by Spectronaut 15 (Biognosys, Schlieren, Switzerland) and the durations of the MS^1^ and DIA scans (TABLE S3) to achieve the recommended average peak sampling of 10 DPPP (6σ) by Pino et al., which corresponds to ∼ 3.25 DPPP (FWHM) calculated by DIA-NN.

As a further optimization of DIA window placement, we introduced a retention time dependency for the window widths in our methods (FIGURE 1). The retention time distribution of tryptic peptides in reverse-chromatography is not completely independent of the m/z. This means that in general peptides with lower m/z values tend to elute earlier as peptides with higher m/z values. In order to use this information to increase the selectivity of the DIA isolation windows, we split the peptide isolation window into 5 ranges of equal size and calculated the precursor m/z distribution from the GPF data for each of these ranges independently. Afterwards the window widths were selected within each range, while the number of windows and so the number of DPPP was kept constant for all ranges. A schematic representation of this strategy in shown in FIGURE 1.

**FIGURE 1:**
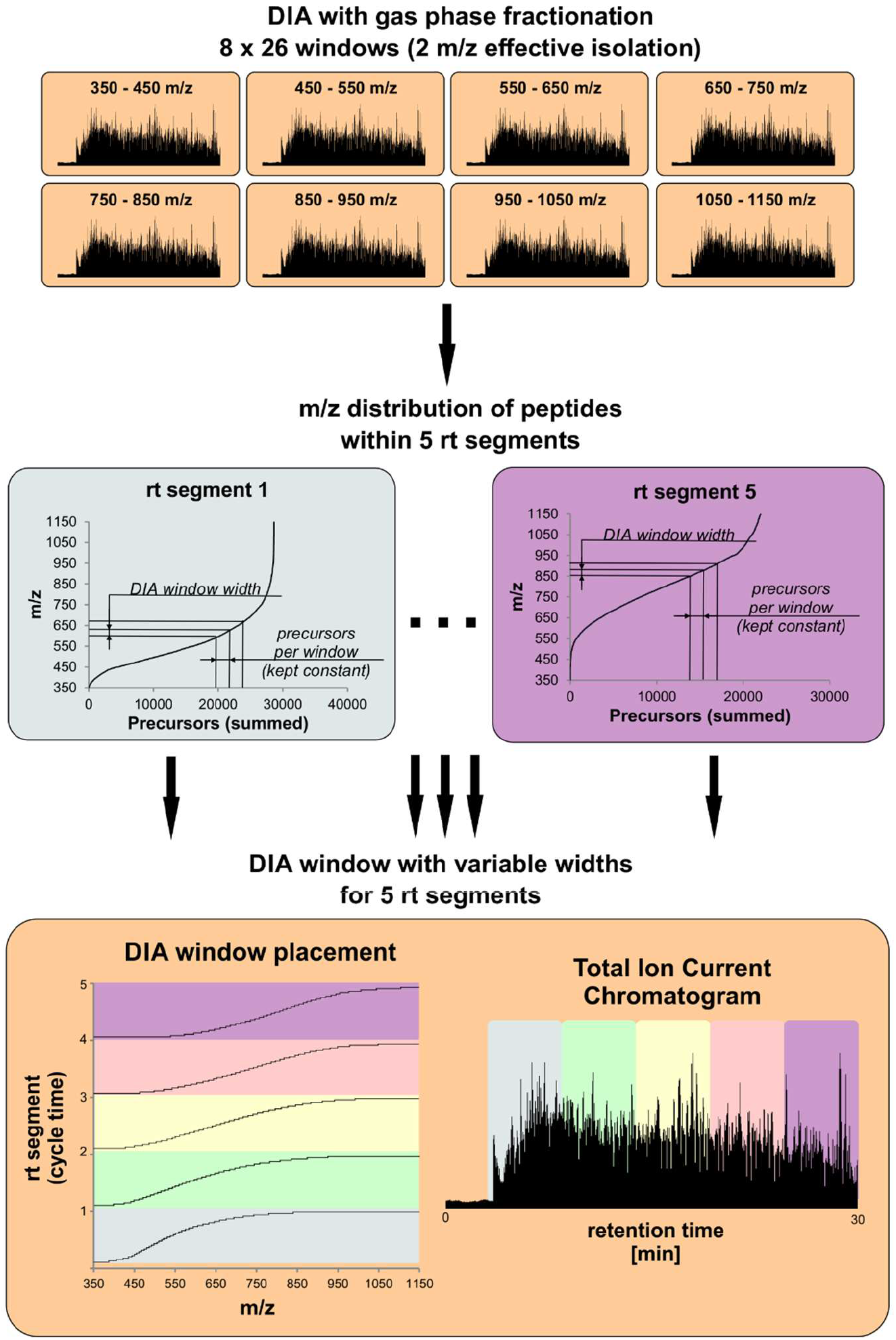
Selection of rt dependent DIA isolation window widths. The m/z distributions of all detectable precursors in HeLa cells within 5 retention time segments were determined from measurements using narrow isolation windows (4 m/z widths with 2 m/z overlap) and gas-phase fractionation (GPF) (8 × 100 m/z, 350 – 1,150 m/z). The resulting m/z distributions illustrate the rt dependency. Isolation window widths were selected separately for each rt segment from the distributions such as the number of precursors identified in the GPF data was kept constant for each window. This information was used to create an instrument method with 5 different rt-dependent isolation schemes but with constant cycle times.

The results of the DIA window optimization experiments are presented in FIGURE 2. All methods were evaluated in triplicate measurements of HeLa cells and independent analysis in DIA-NN using an *in silico* predicted library of the human proteome. Protein identifications increased with decreasing number of DPPP with a maximum of 7,010 proteins using the method with one DPPP (“1 DPPP”, 49 variable windows). However, most proteins (5,956) were consistently quantified with coefficients of variation (CV) below 20% in the “1.25 DPPP” method (39 variable windows), which also led to the highest number of precursor identifications (84,175). Therefore the 1.25 DPPP method was used for evaluation of introducing the retention time dependency of the window widths (5 × 39 variable windows). This strategy led to a slight enhancement of the number of precursor and protein identifications and provided the highest identification numbers in all three categories (87,166 precursors, 7,190 identified proteins, 6,018 proteins quantified with CV < 0.2). In total, the 5 × 39 variable windows method represents an increase of 31% in protein identifications and 56% in precursor identifications compared to the reference method, which was set up according to Pino et al ^17^. It should be noted that this substantial improvement is solely based on optimizing the window acquisition scheme. The influence of lowering the number of DPPP on the FDR of precursor and protein identifications was analyzed by a subsequent analysis of the 5 × 1.25 DPPP and reference data using a combined *in silico* predicted library of the complete human and *E. coli* proteome. FDRs were calculated based on identified *E. coli* sequences at protein and precursor levels (TABLE S9) and were with precursor FDR’s of ∼ 0.1% and protein FDR’s of ∼ 0.2% proven to be very stable across all methods tested. It must be noted, that the number of *E. coli* genes in the database is at the order of 20% of human genes. Although true FDR values are therefore most probably higher, the approach presented nevertheless allows for a relative comparison of the acquisition methods.

**FIGURE 2:**
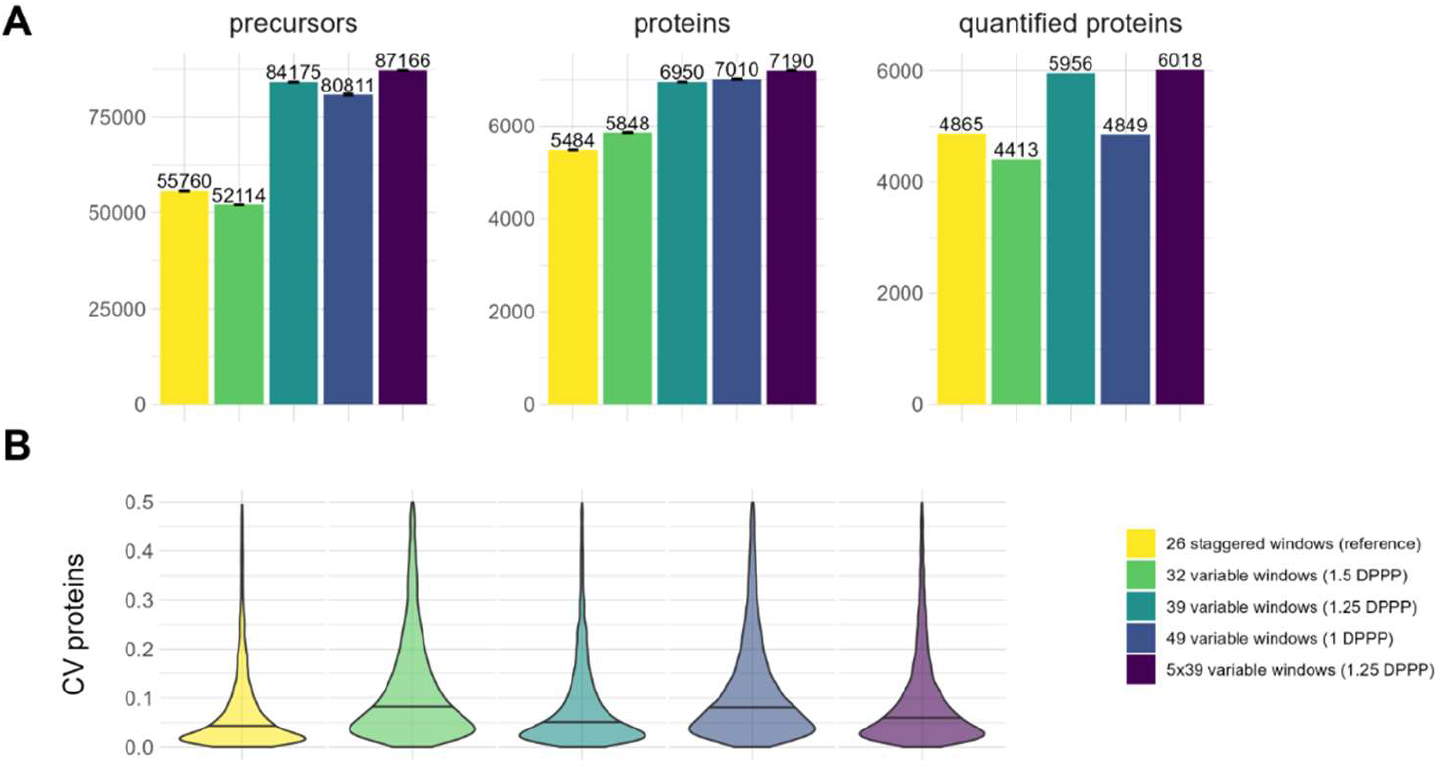
Results of DIA isolation window optimization. HeLa cells were analyzed in triplicates using a 30 min gradient with DIA isolation windows with variable widths. Methods were created for 1, 1.25 and 1.5 data points per peaks (DPPP) on average. Furthermore, a variation of the “1.25 DPPP” method consisting of 5 retention time segments was also included (5×1.5 DPPP). As a reference, 26 staggered windows of fixed width according to Pino et al. were chosen ^17^. Numbers of identified proteins, quantified proteins with coefficients of variation (CV) below 0.2 and identified precursors are compared in A, while violin plots of the protein CV values are shown in B. The median CV values are displayed as black lines.

The protein CV distributions for all acquisition methods are shown in FIGURE 2B. Interestingly, the median protein CV of the 1.25 DPPP method (0.06) exceeds the CV of the reference method (0.05) only slightly, while the 1.25 DPPP method with rt dependent window widths led to another slight increase of the CV (0.07). This is rather unexpected, as the number of DPPP (FWHM) in the reference method is on 2.6 times larger than in the 1.25 DPPP methods. Furthermore, one would expect that the large increase in protein identifications of up to 31% would also contribute to an increase in CV, since it is supposedly mainly “new” low and medium abundant proteins that are identified. In order to analyze this observation in more detail, we extracted peak properties from the data. The results are plotted in FIGURE 3.

**FIGURE 3:**
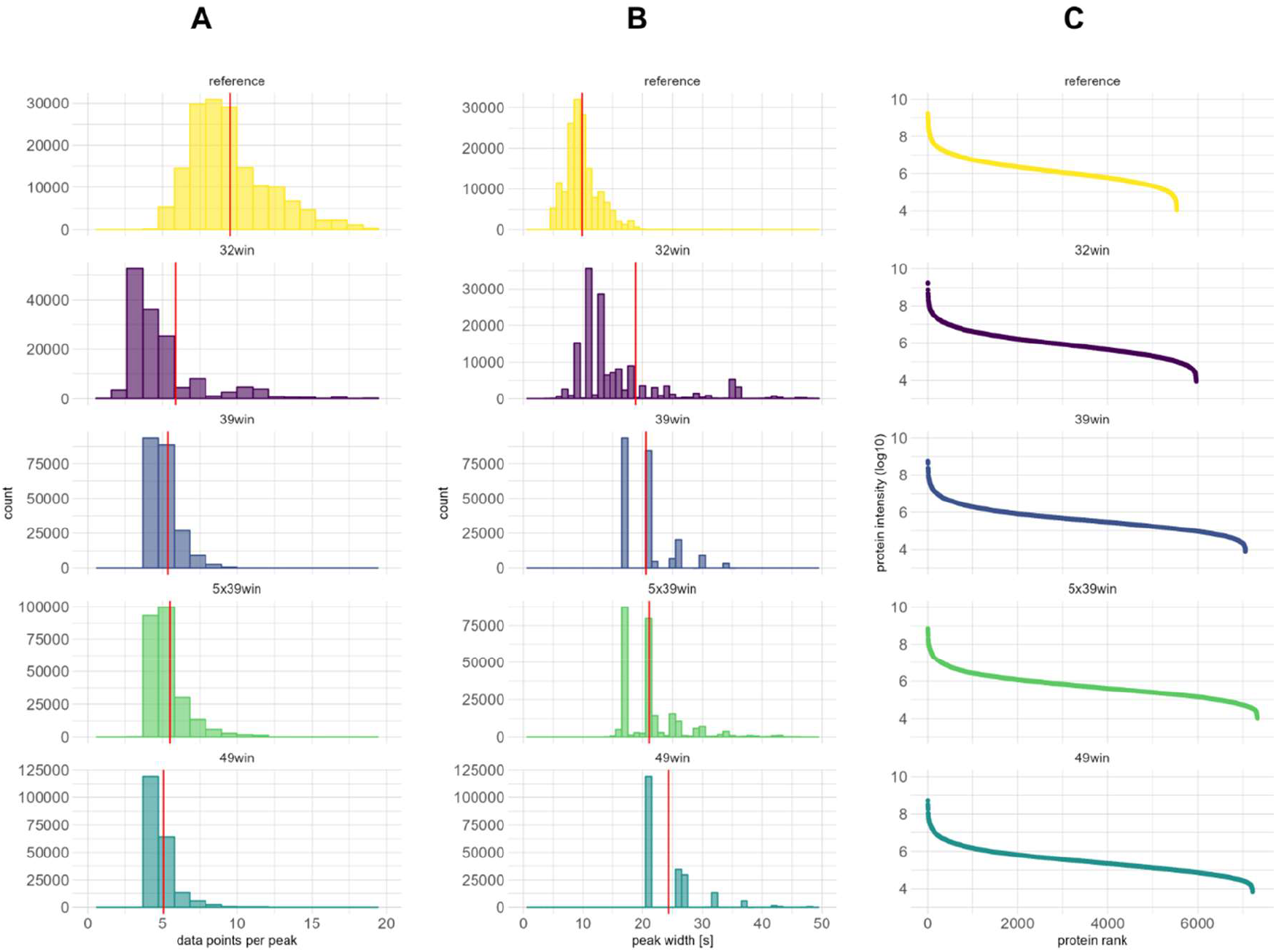
Peak properties of the DIA window optimization data. HeLa cells were analyzed in triplicates using a 30 min gradient with 5 different DIA window isolation schemes. Column A shows the distribution of the data points per peak at base, the red lines represent the mean values. The mean value for the reference is 9.5, which is close to the expected value of 10 used for method creation. The mean value decreases with increasing number of isolation windows (5.9 – 32 windows, 5.4 – 39 windows, 5.5 – 5×39 windows, 5.1 - 49 windows). The histograms in column B represent the LC peak widths of the peptide fragments with the mean values drawn as red lines. The mean value of the reference is 9.8 seconds, which increases with increasing number of isolation windows (18.8 – 32 windows, 20.6 – 39 windows, 21.1 – 5×39 windows, 24.4 - 49 windows). The dynamic range for protein identifications can be read from column C, where the protein intensities (log10) are plotted against the ranked proteins. The dynamic range spans over 5 order of magnitude for all methods.

Column A displays the distributions of the DPPP at base, which is the more commonly used metric in the literature compared to the DPPP at FWHM reported by DIA-NN. The mean value for the reference is 9.5, which is close to the expected value of 10 used for method creation and decreases when the number of isolation windows are increased down to values between 5.9 and 5.1. The distributions of the LC peak widths of the fragment ions display the opposite behavior. The mean peak width of the reference method is 9.8 seconds and increases in conjunction with the number of isolation windows up to values in the range of 18.8 – 24.2 seconds. Please note, that the distributions for the 1.25 and 1.0 DPPP methods become discontinuous due to the long cycle times. As the peptide load and chromatography were unaltered for all methods, the LC peak widths do of course not actually broaden but instead fragment ions could be tracked over a longer period of the LC gradient by DIA-NN. The dynamic range for protein identifications of the methods with varying DPPP’s is shown in column C, where the protein intensities (log10) are plotted against the ranked proteins. The dynamic range spans over 5 order of magnitude for all methods. As the sensitivity for protein identifications does not increase when decreasing the number of DPPP, the increase of the LC peak width must be the result of an improved selectivity and not sensitivity of the precursor identification. To further investigate this, a dilution series (1, 10, 100, 1000 ng) of HeLa samples was analyzed by the 5×1.25 DPPP and the reference method (FIGURE S2). Numbers of precursor and protein identifications are increased for the 5×1.25 DPPP method at 1000, 100 and 10 ng sample load compared to the reference. At 1 ng sample load the numbers are quite comparable. At this low peptide load the number of detectable peaks has decreased so much, that the improved selectivity of the 1.25 DPPP method has lost its advantage for precursor identification. The meaning of the improved selectivity for protein quantification is shown in FIGURE 4, which displays protein features for all proteins consistently identified by all methods

**FIGURE 4:**
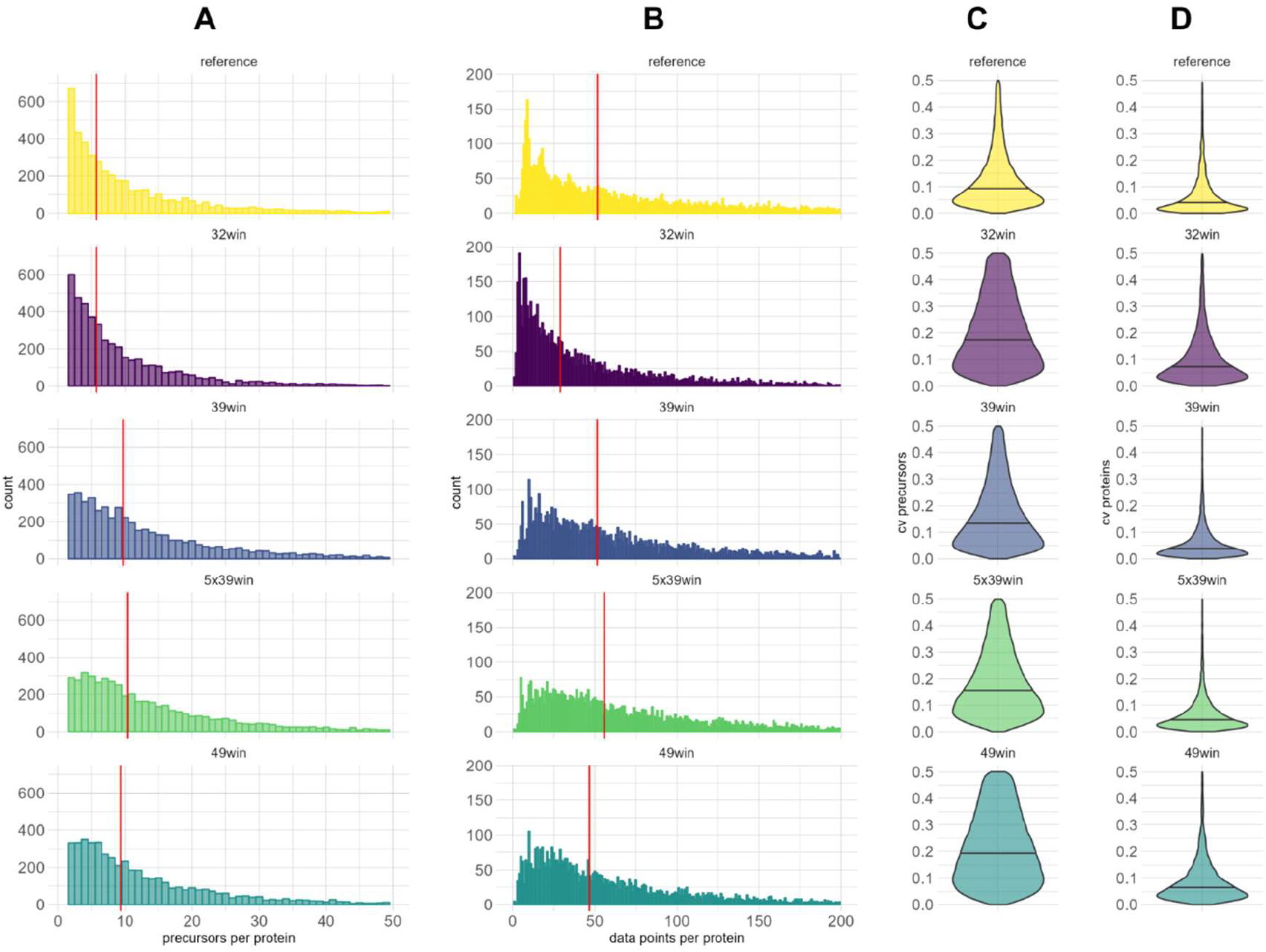
Protein features of the DIA window optimization data. HeLa cells were analyzed in triplicates using a 30 min gradient with 5 different DIA window isolation schemes. The figure represents all proteins, which were consistently identified by all methods (n = 5258). Column A shows the distribution of the number of precursors per protein, with the median values plotted as red lines (5.7 – reference, 5.7 – 32 windows, 9.7 – 39 windows, 10.3 – 5×39 windows, 9.7 - 49 windows). Column B shows the distribution of the data points per protein, with the median values plotted as red lines (55.6 – reference, 29.0 – 32 windows, 51.4 – 39 windows, 51.6 – 5×39 windows, 55.6 - 49 windows). The coefficients of variation of the precursors (C) and proteins (D) are further displayed as violin plots with the median values plotted as black lines.

The number of precursors per protein increases for low DPPP’s compared to the reference from 5.7 (reference) up to 10.3 (5×1.25 DPPP) (FIGURE 4A). If the precursor information is aggregated to proteins, this compensates for the lower number of data points per peak (FIGURE 3A) and so the number of data points per protein are quite similar for the 5×1.25 DPPP and the reference method (51.6 and 55.6). As a result, although the median CV value of the precursors (FIGURE 4C) is clearly lowest for the reference method, the median CV values of the proteins (FIGURE 4D) are very similar for the 5×1.25 DPPP and the reference method (0.044 and 0.038).

This isolation window optimization result contradicts an earlier study performed on the same LC-MS instrument ^18^. In the study of Bruderer et al., reducing the number of DPPP from 11 to 5 had only a small effect on the number of precursor identifications, with 8 DPPP resulted in the best overall performance. Presumably, due to the large reduction of the LC gradient from 120 to 30 min as well as the massive increase of the spectral library (whole proteome predicted vs DDA-based library), selectivity of precursor identification has a much larger role for the overall outcome in this study.

### Classification of the DIA performance with optimized windows

The performance of the optimized DIA method was compared to published data from later generations of mass spectrometers. Therefore, publicly available DIA data of HeLa digests, which were acquired using the 60 samples per day (SPD) method on an EvoSep One LC system coupled either to an Orbitrap Exploris 480 mass spectrometer with and without FAIMS, or a timsTOF Pro instrument were downloaded from PRIDE and compared to the data of this study. All data were analyzed independently for each instrument with enabled MBR using the same *in silico* predicted library of the human proteome in DIA-NN. The results are presented in TABLE 1. Of course, this comparison has severe limitations, as the HeLa digests differ between the studies, the gradient of the 60 SPD method is shorter than the one used in this study (21 vs 30 min) and the peptide loading amount varied between 200 to 1000 ng. Nevertheless, this comparison allows us to get a rough overview to assess the performance of each method when using a whole organism predicted libraryand should therefore help to place the results in a larger context. However, there are several other aspects that need to be considered in more detail. The optimal loading amount for the timsTOF Pro is 200 ng, which is a limitation when analyzing samples of non-limited amount. This issue has been resolved in the recently introduced timsTOF HT, which should therefore further increase the dynamic range of the analysis. The two orbitrap instruments, QE HF and the Exploris 480, have the same scanning speed (12 Hz) at a resolution of 30,000, which was used in this study. The major difference between the two instruments in this experimental setup should therefore be the higher sensitivity of the Exploris 480. However, even when loading 100 ng HeLa samples (FIGURE S2), the number of identified precursors on the QE HF is comparable (42,845) to the data obtained on the Exploris 480 at a sample load of 500 ng (41,479 without FAIMS and 38,756 with FAIMS). This can at least partially be explained by the higher sensitivity of the nanoLC setup used in this study, which uses a lower flow rate in comparison to the Evosep One (300 nL/min vs 1000 nL/min). Using the Evosep One in turn has the advantage of doubling the throughput (60 vs 29 samples per day) with a higher robustness at the same time.

**TABLE 1:**
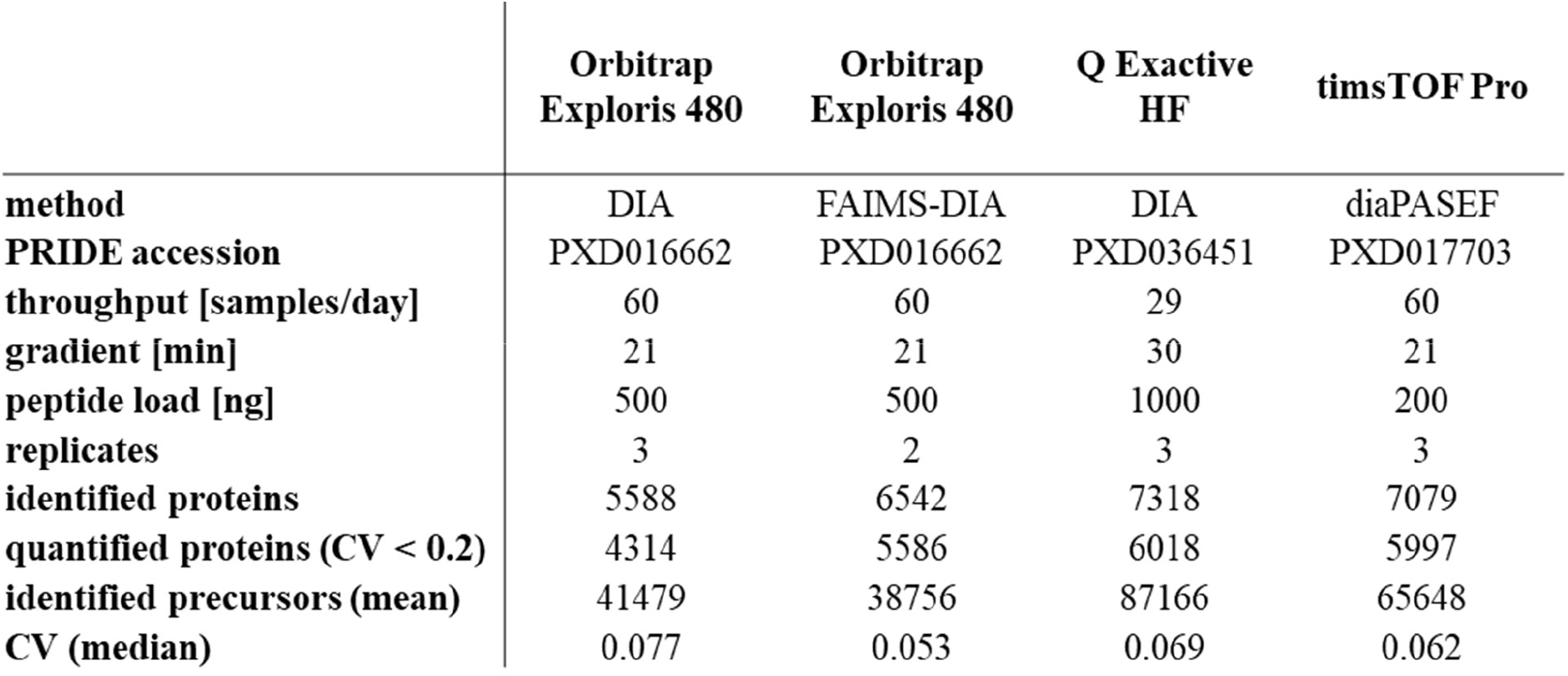
Comparison of short-gradient DIA data of HeLa cells acquired by different MS instruments. Publicly available datasets of short-gradient DIA HeLa data ^1, 2^ were downloaded from PRIDE ^19^ and compared to data from this study acquired using the 5 × 1.25 DPPP method. All data were analyzed using the same *in silico* predicted library of the human proteome in DIA-NN with default settings.

|  | Orbitrap<br>Exploris 480 | Orbitrap<br>Exploris 480 | Q Exactive<br>HF | timsTOF Pro |
| --- | --- | --- | --- | --- |
| **method** | DIA | FAIMS-DIA | DIA | diaPASEF |
| **PRIDE accession** | PXD016662 | PXD016662 | PXD036451 | PXD017703 |
| **throughput [samples/day]** | 60 | 60 | 29 | 60 |
| **gradient [min]** | 21 | 21 | 30 | 21 |
| **peptide load [ng]** | 500 | 500 | 1000 | 200 |
| **replicates** | 3 | 2 | 3 | 3 |
| **identified proteins** | 5588 | 6542 | 7318 | 7079 |
| **quantified proteins (CV < 0.2)** | 4314 | 5586 | 6018 | 5997 |
| **identified precursors (mean)** | 41479 | 38756 | 87166 | 65648 |
| **CV (median)** | 0.077 | 0.053 | 0.069 | 0.062 |

When the DIA-NN results are compared (TABLE 1), it is noticeable that the optimized method of this study provided the highest numbers in all three categories (87,166 precursors, 7,318 identified proteins (overall), 6,018 proteins quantified with CV < 0.2). The median CV’s of all methods are quite similar, except for FAIMS-DIA, which however only represents duplicate measurements. Most strikingly, is the excellent performance of the method optimized in this study with respect to precursor identifications, whose average value e.g. exceeds the data acquired using FAIMS-DIA on an Orbitrap Exploris 480 by 125%. These results show, that the potential of high-throughput DIA-MS is still not fully developed as the strategy for isolation window selection presented in this study could be applied to the latest generation of mass spectrometers as well and that older MS instruments are also well suited for proteomic studies using short gradients.

The proteome depth of the optimized 30 min gradient method was evaluated for cellular proteomes with varying complexity, including *E. coli, S. cerevisiae* and *Homo sapiens* (HeLa). The total protein approach ^20^ was used to calculate the distribution of protein copy numbers in the data (FIGURE 5) and so enable absolute quantification of proteins within a cell. At first, the average protein mass per cell was calculated from total protein content measurements and cell counting of the samples. Afterwards, the copy number of each protein (x) was calculated using the following formula according to Wiśniewski et al. ^20^:

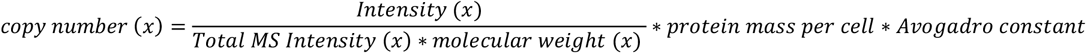

**FIGURE 5:**
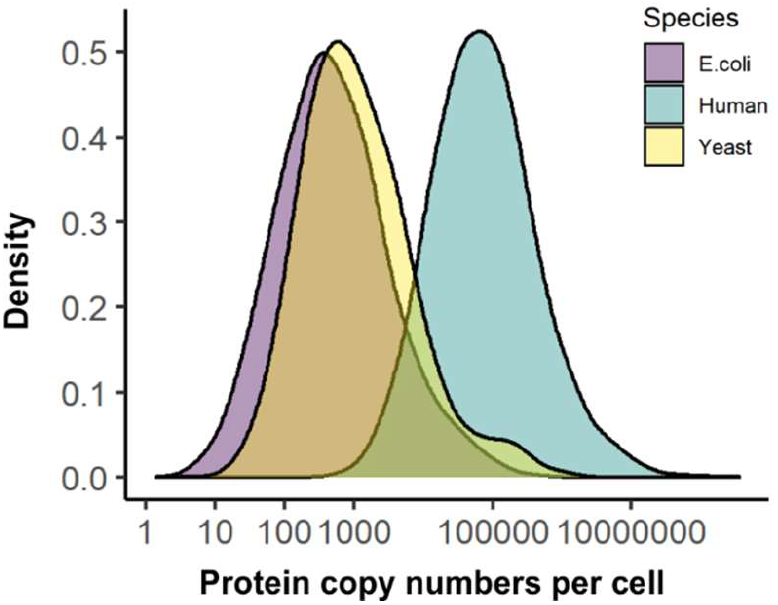
Proteome depth of 30 min gradient DIA data for different species. The density distributions of the protein copy numbers per cell were calculated for HeLa, *S. cerevisiae* and *E. coli* cells using the total protein approach ^20^. The distributions represent all proteins, which were consistently identified in three replicates from a 30 min DIA analysis.

**FIGURE 6:**
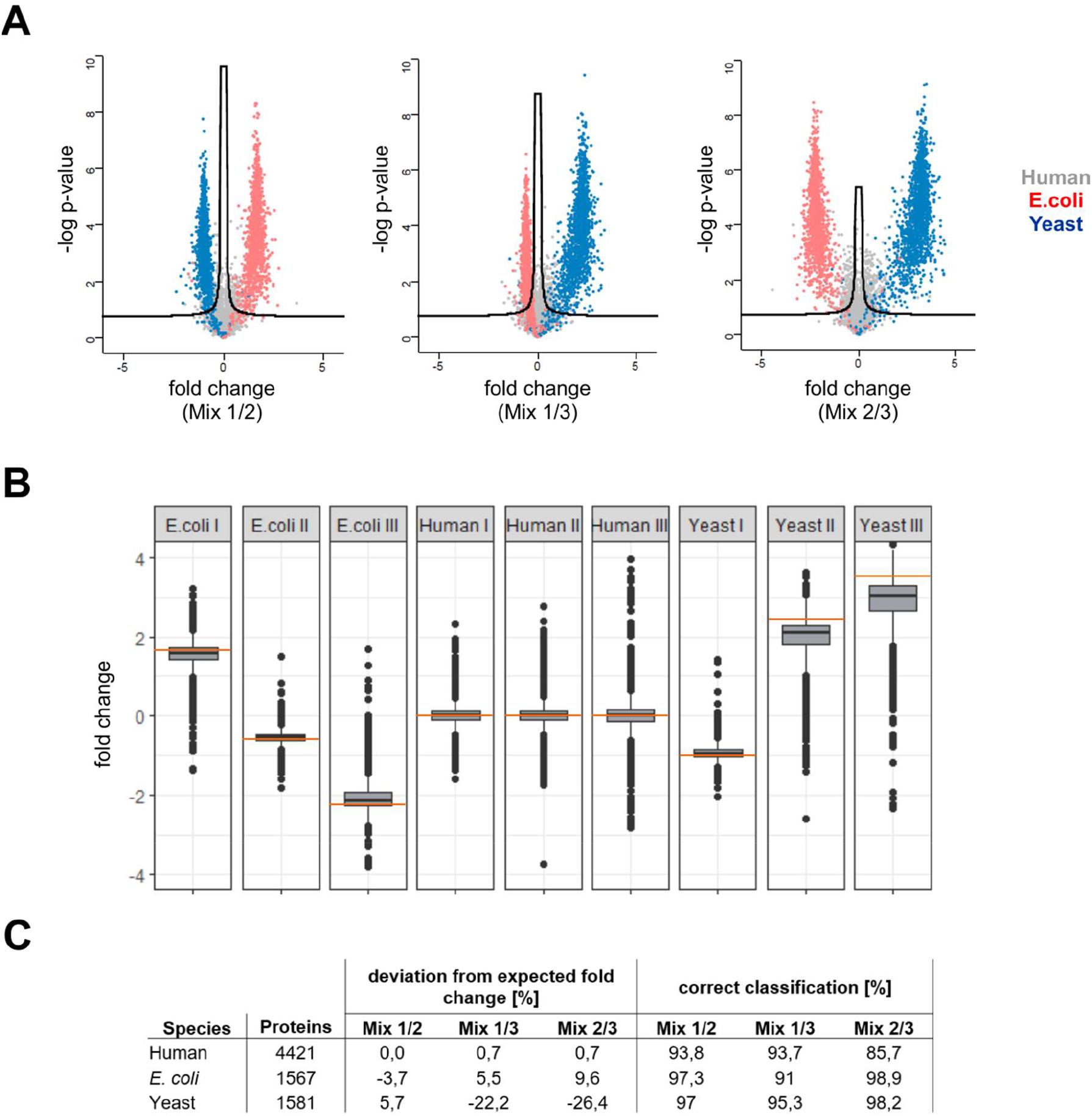
Evaluation of quantification precision and accuracy. Protein quantification accuracy and precision of the 5 × 1.25 DPPP method was analyzed using human, *E. coli* and yeast digests, which were mixed at different ratios in order to obtain three different samples (mix 1-3) corresponding to 9 ratios ranging from −2.2 and 3.3 on a log2-scale (TABLE S). Differentially abundant proteins were detected using t-tests with permutation-based FDR (5%) control. The results of the statistical analysis are visualized as volcano plots (A). The distributions of the observed fold changes are shown as boxplots along with the expected fold change (orange) in B. The deviation of the observed from the expected fold change is summarized in C in conjunction with the number of identified proteins and the ratios of correct classification, which were calculated independently for each species.

Only proteins, which were identified consistently in all three replicates were considered. The observed limits of detection (95th percentile) for the protein copy numbers per cell are 4 for *E. coli* (2549 proteins), 79 for *S. cerevisiae* (4198 proteins) and 5059 for *Homo sapiens* (6801 proteins). This demonstrates, that almost complete bacterial proteomes are detectable in a 30 min gradient, which offers great opportunities for clinical microbiology including antimicrobial resistance detection and species identification as well as for molecular epidemiology of bacterial infections ^21^.

### Evaluation of protein quantification using DIA with optimized windows

The performance of the 1.25 DPPP method with rt dependency was evaluated with respect to protein quantification using mixtures of *Homo sapiens, E. coli* and *S. cerevisiae* digests. Purified peptides were mixed at different ratios in order to obtain three different samples (mixtures 1-3) resulting in 9 different fold changes ranging from 0.2 to 10, which corresponds to a range of −2.2 and 3.3 on a log2-scale (TABLE S1). Differential analysis of protein abundance was performed from triplicate measurements using t-tests with FDR control to account for multiple testing. Percentage of correct protein abundance classification was performed from the statistical results taking the species information of each protein into account. Mixtures were prepared such as the abundance of human proteins was unaltered, while abundance of *E. coli* and *S. cerevisiae* proteins differed between the samples with varying ratios. Volcano plots of the comparisons are shown in figure 5A and the classification results are summarized in 5C. The accuracy of the quantification is visualized as box plots in figure 5B and is summarized as the average percentage deviation from the expected ratio in figure 5C. In general, precision of quantification was high, which resulted in true positive rates > 95% for classification of *E. coli* and yeast proteins in 5 out 6 comparisons. The classification of *E. coli* proteins between the mixtures 1 and 3 was slightly less precise with 91% correct classifications. This comparison has the smallest expected ratio (0.5-fold change), which shows that the precision of classification depends on the abundance ratio. This is quite expectable, as statistical power decreases with lower effect sizes. True negative rates of the human proteins was 94% in two comparisons, which reflects the FDR of 5% used for statistical testing quite well, but decreased to 86% when analyzing mixtures 2 and 3. This comparison contains high expected ratios for *E. coli* (−5-fold change) and *S. cerevisiae* (10-fold change) proteins, which might have introduced some challenges for normalization and statistical testing, which could not be addressed by the quite simple strategy used in this study. The error of abundance ratio estimation was below 10% in 7 out of 9 comparisons and increased up to 25% for yeast proteins when large alterations are expected (−5 and 10-fold change). These results are highly encouraging and show that the short-gradient DIA method with low number of DPPP is well suited for differential protein expression analysis.

## CONCLUSION

High-throughput proteomics benefits greatly from advancements of short gradient DIA methodologies. In this study, we demonstrate that optimization of data acquisition even without any hardware adaptation has a huge untapped potential to increase proteome depth. The strategy presented enabled us to acquire proteomes using a rather old mass spectrometer up to a depth and precision, achievable to date only by later generation of instruments. Our findings should spread the availability of high-throughput proteomics platforms further as it proves, that actually many older instruments can be effectively used for this task. The data also shows, that almost complete bacterial proteomes can be analyzed in just 30 min gradient time. This opens up great opportunities for microbiology applications, a discipline with huge lacks of knowledge on proteome level because of the limited availability of antibodies. The analysis of complete bacterial proteomes in high-throughput mode is thought to be helpful to fully uncover the diagnostic potential of proteomics in clinical microbiology and to provide deeper insights into bacterial evolution when proteomics is used to complement genomics in molecular epidemiology. Furthermore, the presented optimization strategy should in principle be applicable also to faster scanning mass spectrometers and could thus enable further improvements of proteome depth in short gradient DIA proteomics. However, it should be noted, that the proposed strategy is only intended to be used for analyzing protein-level data and not peptide-level data, e.g. for studying post-translational modifications or proteoforms.

## Access to proteomics data

The mass spectrometry proteomics data have been deposited to the ProteomeXchange Consortium (http://proteomecentral.proteomexchange.org) via the PRIDE partner repository with the dataset identifier PXD036451 ^19^.

## CONTRIBUTIONS

J.D. and P.L. conceptualized and designed the study. J.D. and A.S. performed the experiments.

J.D. analyzed the data, prepared figures and wrote the initial draft of the manuscript. C.B. assisted with data analysis. All co-authors contributed to writing, editing, and reviewing the manuscript.

## SUPPORTING INFORMATION

The following supporting information is available free of charge at ACS website http://pubs.acs.org

### Supporting tables

Table S1: Species mix preparation

Table S2: Isolation windows for 8 gas-phase fractions with overlapping windows

Table S3: Number of isolation window calculations

Table S4: Isolation windows corresponding to 1.5 data points per peak (FWHM, 32 windows)

Table S5: Isolation windows corresponding to 1.25 data points per peak (FWHM, 39 windows)

Table S6: Isolation windows corresponding to 1.0 data points per peak (FWHM, 49 windows)

Table S7: Isolation windows corresponding to 10 data points per peak (calculated according to Pino et al.)

Table S8: Isolation windows with retention time dependancy corresponding to 1.25 data points per peak (FWHM, 5 × 39 windows)

Table S9: False discovery rate comparison in HeLa samples using a combined library of the human and *E. coli* proteome

### Supporting figures

Figure S1: DIA with retention time dependency method setup on a Q Exactive

