## Supplementary material for "Increasing proteome depth while maintaining quantitative precision in short gradient data-independent acquisition proteomics": FIGURE S

**TABLE OF CONTENTS**

1. **Supplementary figures**

**Figure S1:** DIA with rt dependency method setup on a Q Exactive ……….…	3

**Figure S2:** Results of HeLa dilution series DIA measurements ……….……..	4

2. **Supplementary tables (SI Tables.xlsx)**

**Table S1:** Species mix preparation

**Table S2:** Isolation windows for 8 gas-phase fractions with overlapping windows

**Table S3:** Number of isolation window calculations

**Table S4:** Isolation windows corresponding to 1.5 data points per peak (FWHM, 32 windows)

**Table S5:** Isolation windows corresponding to 1.25 data points per peak (FWHM, 39 windows)

**Table S6:** Isolation windows corresponding to 1.0 data points per peak (FWHM, 49 windows)

**Table S7:** Isolation windows corresponding to 10 data points per peak (calculated according to Pino et al.)

**Table S8:** Isolation windows with retention time dependancy corresponding to 1.25 data points per	peak (FWHM, 5 × 39 windows)

**Table S9:** False discovery rate comparison in HeLa samples using a combined library of the human and *E. coli* proteome

centre of window #33

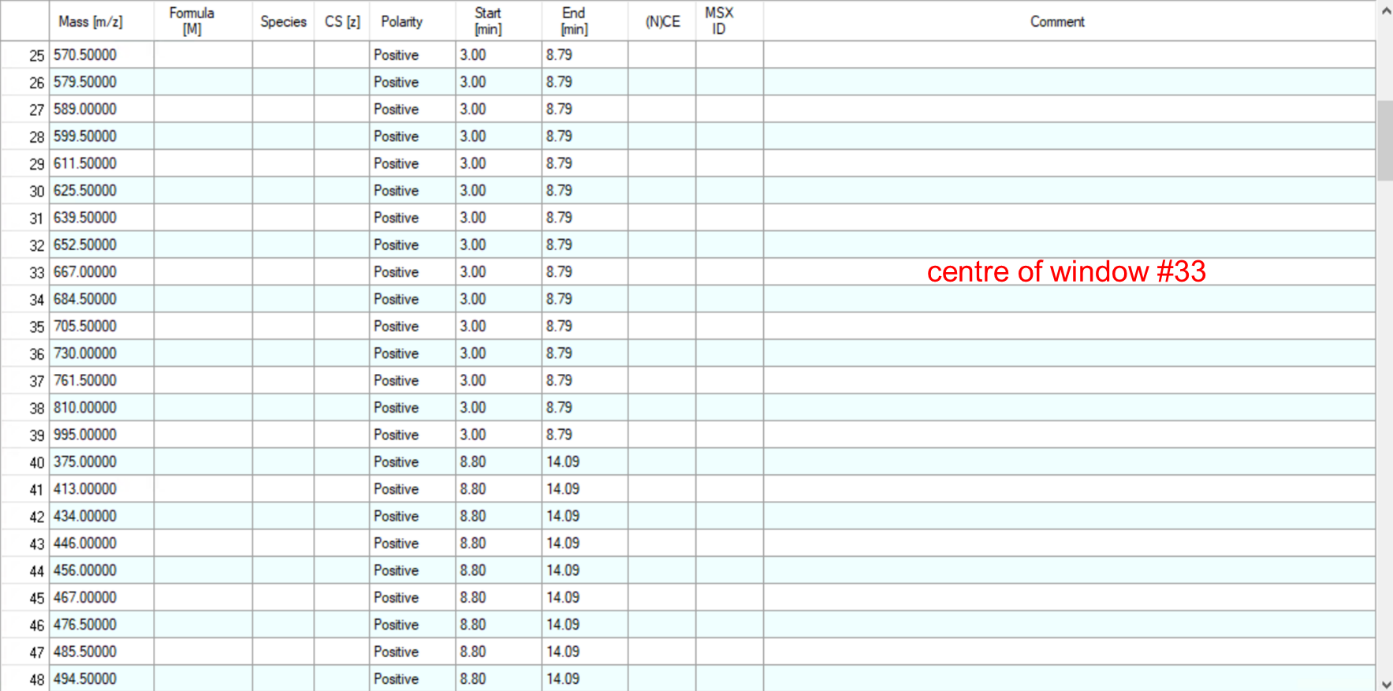

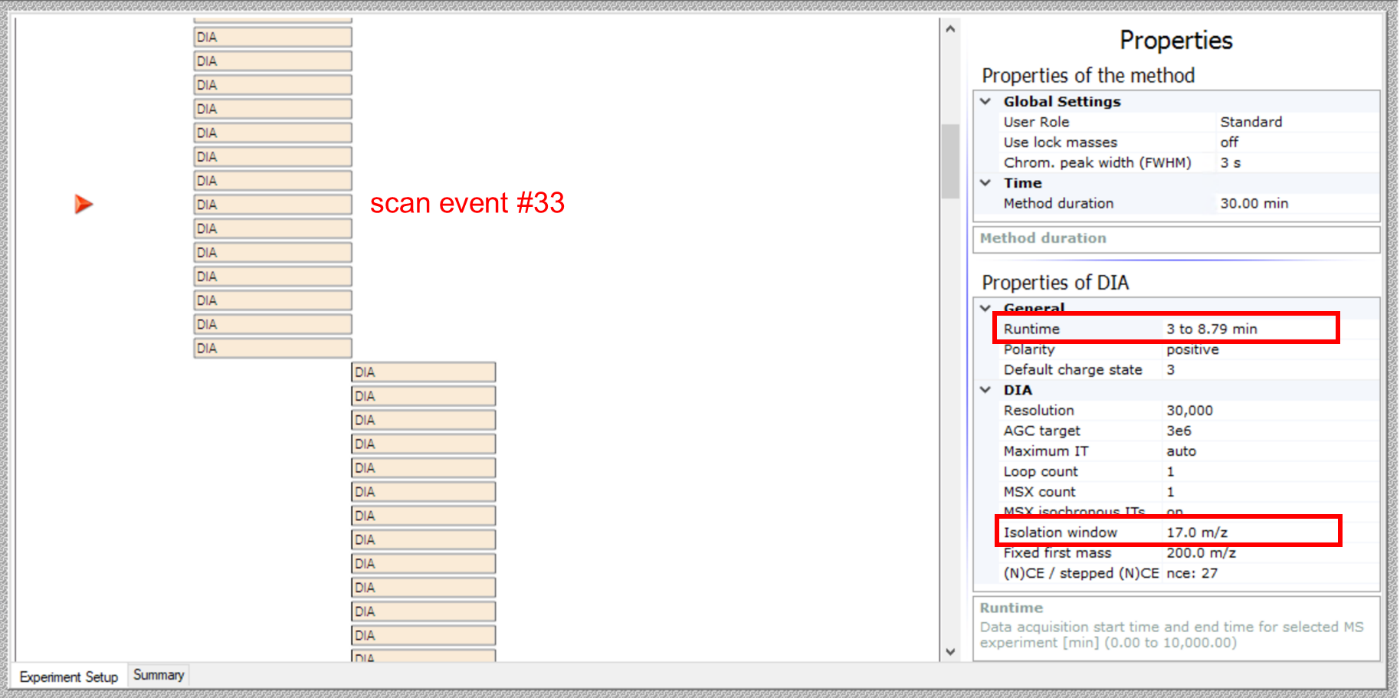

**Figure S1:** **DIA with retention time dependency method setup on a Q Exactive.**

In order to introduce retention time dependent segments of DIA cycles with fixed cycle times but varying window widths, the window centers are deposited in the inclusion list of the QE method editor along with start and end times. Afterwards, the runtime and the isolation window width of each single DIA scan event need to be set in accordance to the order of the inclusion list. In total the 5 x 1.25 DPPP method consists of 1 x MS1 scan event (run time = 3 to 30 min) and 195 DIA scan events.

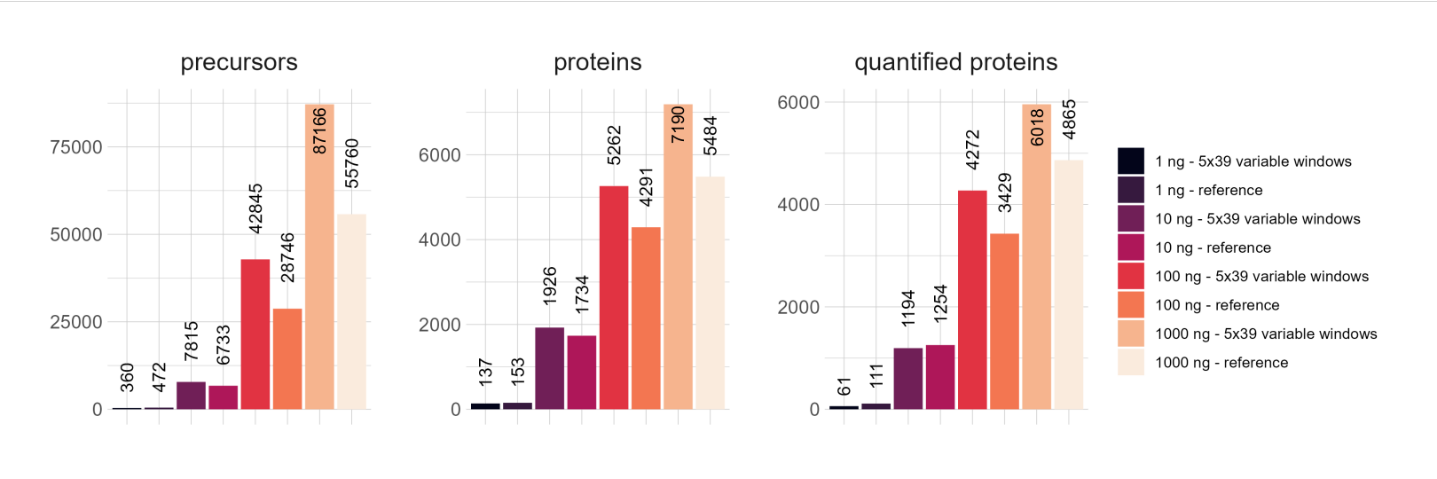

**Figure S2:** **Results of HeLa dilution series DIA measurements**

HeLa samples were diluted (1000, 100, 10, 1 ng peptide load) and analyzed in triplicates using a 30 min gradient with 5 x 39 DIA isolation windows with variable widths (5x1.25 DPPP). As a reference, 26 staggered windows of fixed width according to Pino et al. were chosen. Mean values of the numbers of identified proteins, quantified proteins with coefficients of variation (CV) below 0.2 and identified precursors are displayed.
